# Persistence of FoxJ1^+^ Pax6^+^ Sox2^+^ ependymal cells throughout life in the human spinal cord

**DOI:** 10.1101/2022.08.01.502176

**Authors:** Chantal Ripoll, Gaetan Poulen, Robert Chevreau, Nicolas Lonjon, Florence Vachiery-Lahaye, Luc Bauchet, Jean-Philippe Hugnot

## Abstract

Spinal cord ependymal cells have stem cell properties in mice. They surround the central canal and keep expressing spinal cord developmental transcription factors. Similar cells exist in young humans however their persistence with aging is debated.

We clarified this issue by collecting 17 spinal cords from organ donors, aged between 37 and 83 years old. We examined the presence of ependymal cells using immunohistochemistry on lightly-fixed tissue.

We found the presence of cells expressing the typical ependymal marker FOXJ1 in the spinal cord central region in 100% of cases. In addition, a lumen surrounded by FOXJ1+ cells was observed in half of the cases. Like in mice, these human ependymal cells maintain the expression of SOX2 and PAX6 proteins together with RFX2 a master transcriptional regulator of ciliogenesis and ARL13B, a regulatory GTPase enriched in cilia. Reminiscent of the situation observed in mice and in young human spinal cord, a fetal-like regionalization of neurodevelopmental transcription factors was observed in three donors aged over 75 years: MSX1 and ARX/FOXA2 was preferentially expressed by dorsal and ventral ependymal cells, respectively.

These results provide new evidence for the persistence of ependymal cells expressing neurodevelopmental genes throughout human life. The persistence of these cells in humans opens new opportunities to regenerate the spinal cord.

## Introduction

The spinal cord is affected in many central nervous system diseases such as traumatic injuries, amyotrophic lateral sclerosis or multiple sclerosis for which very few curative treatments exist. Some animals like Urodela are able to regenerate their spinal cord after lesion, including motor neurons and oligodendrocytes^1^. This fascinating phenomenon is based on the presence of a pool of stem cells, resembling fetal neuroepithelial cells and persisting around the central canal of the adult spinal cord, the so-called ependymal region or ependyma. After lesion, these ependymal cells activate, proliferate and then differentiate into neurons and glial cells^1–3^. Similar ependymal cells with stem cell properties exist in the adult spinal cord in mammals, notably in mice. These cells are also located around the central canal constituting a stem cell niche^4–6^. They keep expressing several spinal cord developmental transcription factors such as Arx, Msx1, Pax6, Sox2,4,6,11^7^. In particular, we observed that the niche has an embryonic-like dorsal-ventral regionalization and we demonstrated the presence of specific radial cells in the dorsal and ventral parts. These radial cells are remnants of the embryonic spinal cord roof and floor plates which are well-known developmental organizing centres^8^. In culture, a fraction of these ependymal cells can generate passageable neurospheres (i.e. clonal expansion of stem and progenitor cells) which when placed in differentiation conditions generate mainly astrocytes and few oligodendrocytes and neurons^9,10^.

Theoretically, such multipotent ependymal cells showing a high capacity for proliferation could be used to alleviate spinal lesions if one was able to control their fate. However one difficulty in harnessing spinal cord ependymal cell potential is that most of our knowledge on these cells is based on rodents. Rodents are short-lived animals which separated from primates at least 100 million years ago. It is thus important to study spinal cord ependymal cells in humans who have a much longer lifespan and as major differences may be present. Unfortunately there are only few works done on spinal cord ependymal cells derived from human tissues, even less using adult tissues. One main obstacle is to get access to well-preserved human tissue to perform histology, cell isolation and culture.

Very few groups have addressed the persistence of spinal cord ependymal cells with aging and the results are somewhat divergent as it is often the case when dealing with human tissues. One group reported in 2015 that the human central canal region disappears throughout the spinal cord from early childhood in the majority of individuals, being substituted by astrocyte gliosis and cells resembling ependymoma (tumors arising from the ependymal region)^11^. No Sox2^+^ or CD133^+^ (PROM1) cells, two markers of ependymal cells, were found after the second decade. In contrast, in 2019, we reported a detailed work done on two donors aged seventeen and forty-six years old, where we observed a typical ependymal region with a lumen^7^. RNA profiling of these cells combined with immunofluorescence validation showed a very good conservation of this region between Humans and mice (i.e conserved expression of 1,200 genes including 120 transcription factors). Human ependymal cells keep expressing markers for immaturity such as nestin, vimentin and several spinal cord developmental transcription factors (Arx, Msx1, Meis2, Sox2/4/6/11). They also express the very specific marker for ependymal cells, namely FoxJ1.

This discrepancy prompted us to address further the loss or persistence of ependymal cells in the human spinal cord during aging by collecting seventeen new spinal cords from organ donors and examining the presence and organization of these cells using immunochemistry.

## Materials and Methods

### Human spinal cord extraction

The human spinal cords were dissected at the Montpellier Hospital from organ donors under the approval of the French institution for organ transplantation (Agence de Biomédecine). An informed consent from the families was obtained by the organ procurement organization for this study. Before vascular aortic clamping, blood was replaced by perfusion with 4°C-chilled IGL1 (Institut Georges Lopez, Lissieu, France) or Custodiol (Essential Pharmaceuticals, Durham, UK) organ preservation solutions for abdominal or thoracic organ collection, respectively. The duration between vascular clamping and spinal cord extraction was of four hours on average, when heart, lungs, liver, and kidneys were removed, and an average of two hours when only the liver and kidneys were removed. Body cavities were cooled with crushed ice. Surgery was performed as described previously^12^ and the thoracic or lumbar segments were immediately placed on ice before processing for immunohistochemistry. Donor characteristics and elapsed time between brain death, clamping and spinal cord extraction are presented in Table S1.

### Histology

Spinal cord segments were fast frozen in a 40 ml flat-bottom plastic tube immersed at −80°C in a pre-chilled isopentane bath placed in a SnapFrost® apparatus (Excilone, Elancourt, France) and then stored in a −80°C freezer. Spinal cords were cryosectioned transversally (14µm thick sections) using a NX70 cryostat (Microm Microtech, Brignais, France) and sections were placed on *SuperFrost*® Plus Menzel Gläser slides (Thermo Scientific, Illkirch, France). Slides were stored in a - 80°C freezer until used. Before immunolabeling, sections were lightly fixed in a 4% paraformaldehyde-PBS (phosphate buffer saline) solution for 20 minutes at 4°C and then washed twice with PBS. Antibodies and dilutions used for stainings are listed in Table S2. Immunohistochemistry was performed using polink-2 HRP Plus rabbit/mouse/goat DAB detection kits (Diagomics, Blagnac, France). Primary antibodies were applied on sections overnight. Images were taken with a Nikon Eclipse microscope. To evaluate the presence of a closed vs open central canal, sections were stained with hematoxylin staining. The number of examined sections is indicated in Table S3.

## Results

To probe the presence of ependymal cells during aging, we collected spinal cords from seventeen organ donors (thoracic or/and lumbar levels) aged between 37-83 years (donors are described in Table S1). The spinal cord was removed within two to four hours after aortic clamping in most cases and six to twenty-seven hours after death. In order to preserve antigens and avoid tissue shrinkage due to formaldehyde fixation^13^, the spinal cord was frozen without fixation and IHC was done on lightly-fixed sections to preserve antigens. We started our analysis by examining the expression of the transcription factor FOXJ1 which is a very good marker of ependymal cells in the mouse and human spinal cord^14^. Fig 1A illustrates the presence of FOXJ1+ cells around the central lumen of two 77 and 83 y. old donors. FOXJ1+ cells were also detected in all other examined cases (13/13) (fig.1B, all cases are presented on fig. S1). We noted that in approximately 50% of spinal cords, a patent lumen was not observed and FOXJ1+ cells rather appeared as a mass of cells located in the central region (see for instance donor 12, upper thoracic, fig S1). When a central canal was present we also observed that in addition to FOXJ1^+^ cells surrounding the lumen, isolated FOXJ1^+^ cells were often found away from the lumen (red arrows on fig. 1). In addition to FOXJ1, in mice and in young human, ependymal cells also express SOX2, a well-known neural stem cell transcription factor together with PAX6, a homeodomain transcription factor which is highly expressed by spinal cord neuroepithelial cells during development^7^. As illustrated in fig 1A, SOX2 and PAX6 patent stainings were present in cells surrounding the lumen, of both 77 and 80 y. old donors, as well as all other examined cases (9/9, fig. 1B). We previously reported a specific expression of cilia-related genes by human and mouse ependymal cells^7^. To validate this expression in aged samples, we performed IHC for RFX2, a master transcription factor controlling ciliogenesis^15^ and ARL13B, a regulatory GTPase highly enriched in cilia^16^. Strong nuclear expression of RFX2 and ARL13B stainings at the border of ependymal cells were observed in all examined cases (fig. 2A, B, C). Altogether these results demonstrate the persistence of ependymal cells expressing ARL13B, FOXJ1, PAX6, RFX2, SOX2 in the human spinal cord during aging.

**Figure 1:**
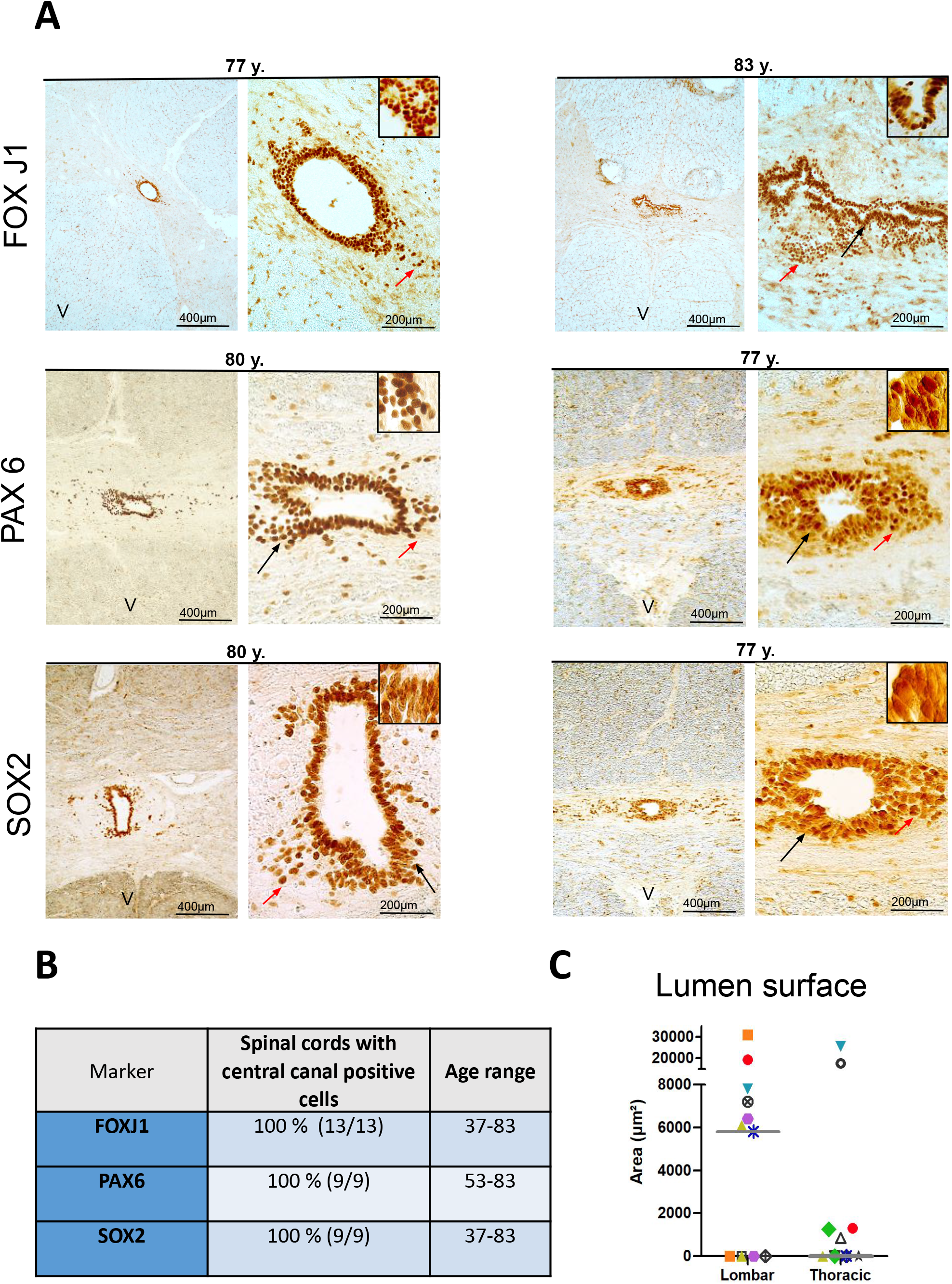
Detection of ependymal cells in aged spinal cords. ***A***. Immunohistochemistry for indicated proteins. Left-hand images are low magnification images (x25) showing the specific expression of explored markers by cells around the central canal. Black arrowhead indicates the magnified zone presented in the inset. Red arrows show stained cells away from the central canal.The **V** letter on images indicates the ventral part of the spinal cord. ***B:*** summary of staining results. ***C:*** Scatter plot of the lumen surfaces measured in lumbar and thoracic spinal cords. Each donor is depicted by a different coloured symbol. Grey bar shows the median value.

**Figure 2:**
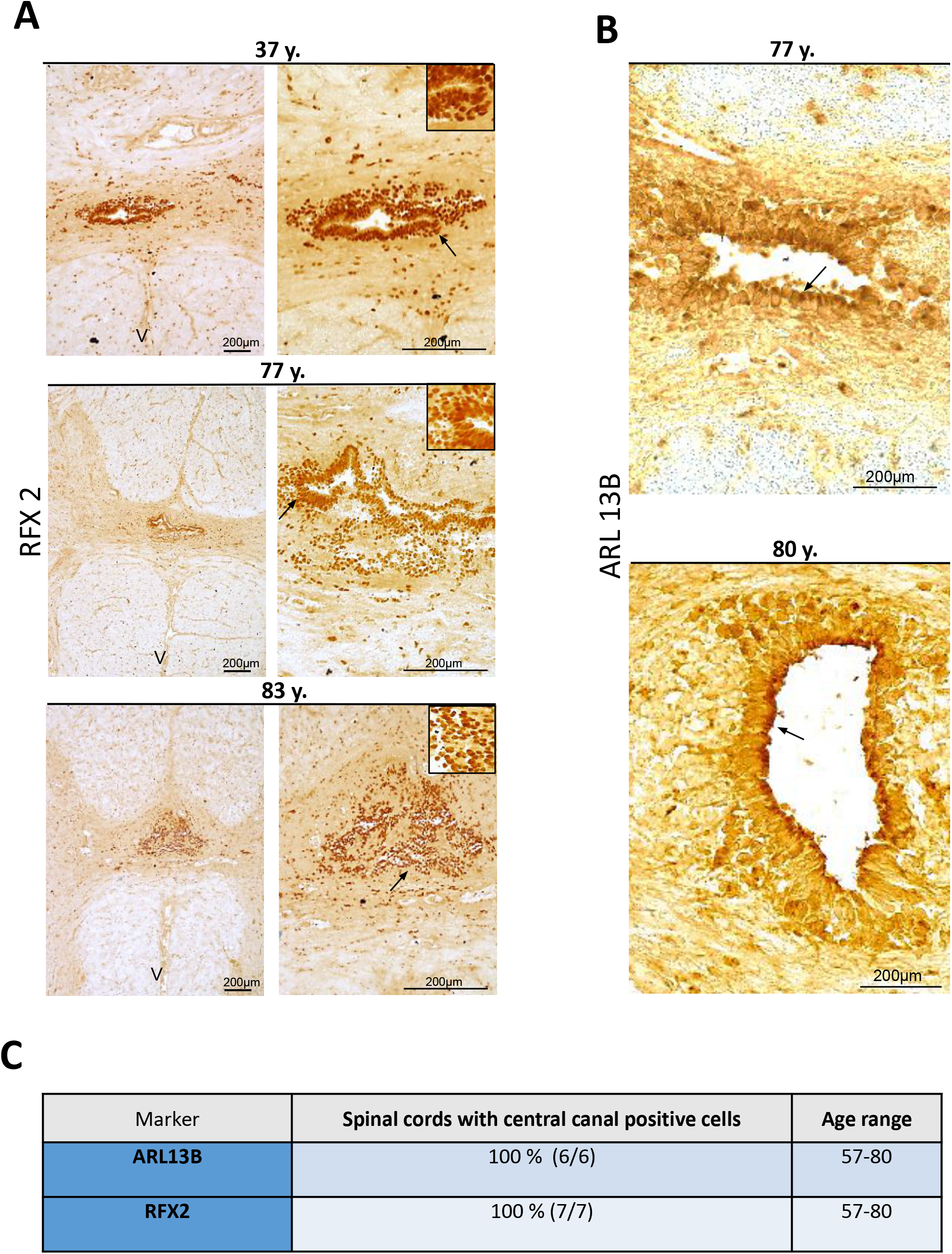
Detection of cilia-associated proteins in aged spinal cords. ***A, B*** Immunohistochemistry for indicated proteins. ***A***: Stainings for RFX2. Left-hand images are low magnification images (x25) showing the specific expression of RFX2 by cells around the central canal. Black arrows indicate the magnified zone presented in the inset. The **V** letter on images indicates the ventral part of the spinal cord. ***B***: Stainings for ARL13B. Arrows show accumulation of staining at the apical side of ependymal cells. ***C:*** summary of staining results.

We previously reported a regionalized expression of spinal cord developmental transcription factors, namely ARX/FOXA2/MSX1, in the mouse and human ependyma^7^. This pattern of expression is reminiscent of the situation during spinal cord development where MSX1 is expressed by cells in the dorsal neuroepithelium while ARX and FOXA2 expression are restricted to the ventral neural tube^8,17^. We performed IHC for these 3 factors to see whether this fetal-like regionalization could also be maintained with aging. As presented in fig. 3, for the 3 studied cases, we observed a weak expression of MSX1 by most ependymal cells however a group of dorsal cells clearly showed a higher MSX1 staining. Conversely to MSX1, ARX and FOXA2 stainings were confined to a group of cells located in the ventral part of the ependyma (fig. 3).

**Figure 3:**
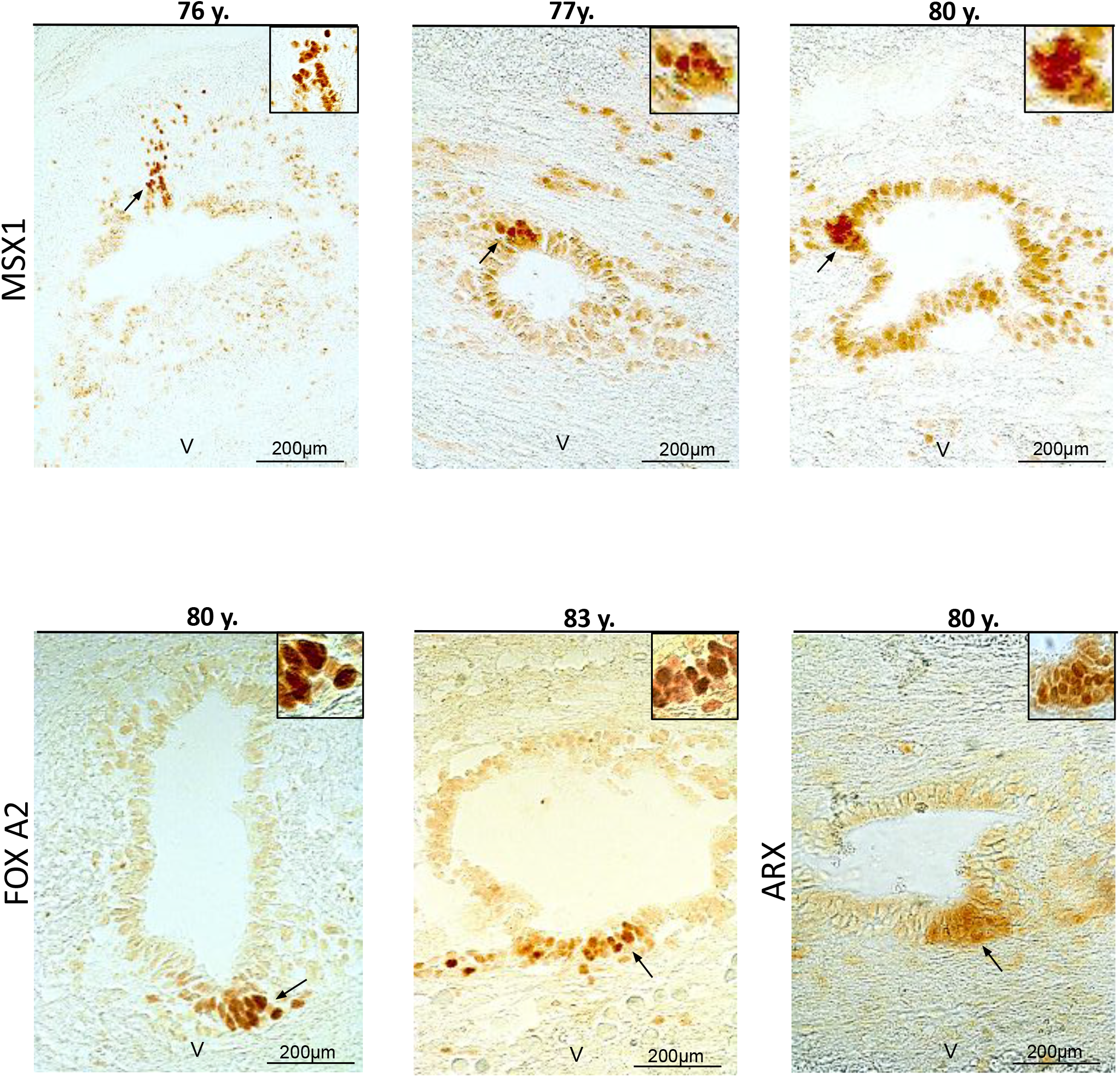
Maintenance of a dorsal-ventral regionalization of the central canal in aged spinal cords. Immunohistochemistry for indicated proteins in three aged donors. Black arrows indicate the magnified zone presented in the inset. The **V** letter on images indicates the ventral part of the spinal cord.

Finally, a loss of central canal and stenosis have been previously reported during aging of the human spinal cord^11,18–20^. We found three types of situations in our cohort (Table S4): **1-**cases in which all examined sections had a lumen (45 % of explored segments). It is worth noting that this lumen was not always circular and could be distorted (see for instance fig.1A top right-hand image, 83 y. old donor). However a wide and round lumen could be observed even in old donors (fig.1A left-hand image). The surface of the lumen is generally larger in the lumbar spinal cord than in the thoracic segment (fig. 1C) where the lumen can be very small (see for instance fig.1A PAX6 staining, 77 y. old donor). **2**-cases in which no patent lumen was found (40 % of explored segments). Examples for this type of pattern are presented in fig. S2A. **3**-cases in which, depending on sections in the same segment, we noted the absence or the presence of a central canal (15 % of explored segments, fig. S2B). To explore further this type of pattern, we performed serial sectioning of 2 samples over a 2 and 5 mm length, respectively. Images presented in fig. 4 showed that there was an alternation of open and closed central canal in the first donor. As for the second examined donor, all sections over 5 mm were closed (not shown).

**Figure 4:**
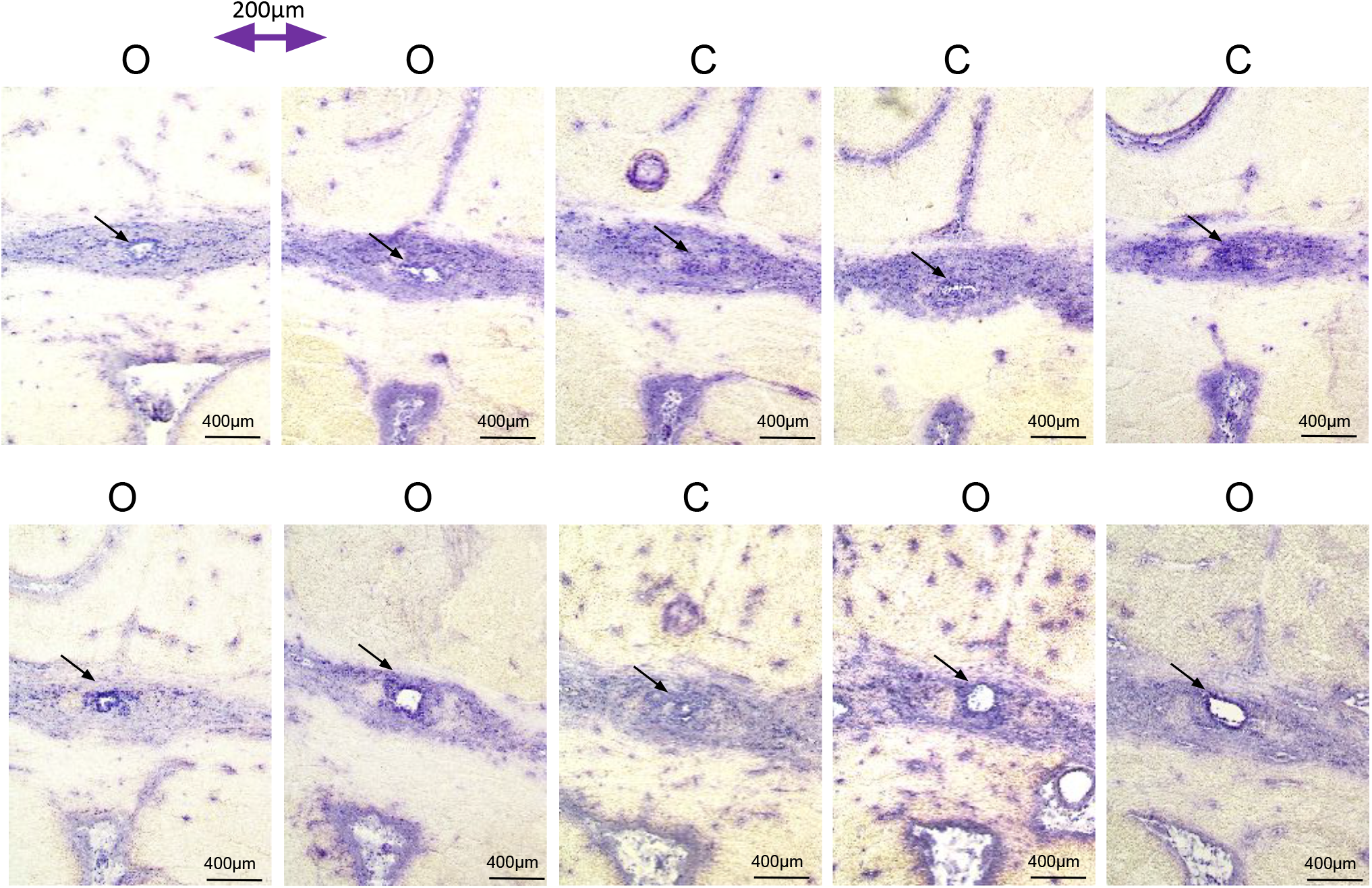
Alternation of closed-open central canals. The series of images shows hematoxylin-stained sections, spaced at 200 µm each, from a 77 y. old thoracic spinal cord. The O and C letters indicate open and closed central canal, respectively.

### Discussion

In this article, we reported that ependymal cells expressing FOXJ1 and spinal cord neurodevelopmental and stem cell proteins (ARX, FOXA2, MSX1, PAX6 and SOX2) are maintained during aging of the human spinal cord. These results are in contrast to previous results suggesting that these cells are lost with aging and that SOX2 expression is no longer detected^11^. Several technical considerations may explain this discrepancy. First, we used organ donors as a source of spinal cords. Compared to autopsy, generally used in most studies on human spinal cord ependyma, organ donors were perfused with an organ preservation solution and the spinal cord was removed only a few hours after clamping, typically two to four hours. This procedure is likely to preserve ependymal cells as close as possible to their original state. Second, after collection, the spinal cord was directly frozen at low temperature without fixation and IHC was performed on frozen sections which were only lightly fixed. In contrast, most studies on the human spinal cord ependyma are based on an extended paraformaldehyde fixation followed by paraffin embedding which have been shown to remove lipids and induce important tissue shrinkage (up to 60% for mouse brain^21^). It is possible that this histological procedure affects the structure of the ependymal region and the reactivity of ependymal cells to antibodies.

Within our cohort, we found that around 50% of cases showed a disorganized ependyma with no obvious lumen. This is consistent with previous works reporting partial or complete stenosis of the ependyma with aging^11,18–20^. However it is important to note that even in this situation, cells expressing FOXJ1, PAX6 and SOX2 were still detected (fig. S2A). We could observe an epithelial organization surrounding a lumen in approximately 40% of spinal cords. We also observed cases containing both open and closed central canal regions, suggesting that disorganization of the central canal region is not homogenous along the spinal cord. In fact, by examining 232 cases, Milhorat, et al. found only four entire occlusions (1.7%) of the entire central canal along the seven levels examined^19^. Combined with our observation of closed and open ependyma regions in the same donor, these data suggest that obliteration of the central canal is discontinued.

We previously reported that in mice and humans, aged 17 and 46 y. old, the ependyma shows an embryonic-like organization with a differential dorsal-ventral expression of spinal cord neurodevelopmental transcription factors such as ARX/FOXA2/MSX1. This organization very likely arises from the persistence of cells originating from the neural tube roof and floor plates. Strikingly, we found that such an organization can be observed even in donors over 76 years old. During development, these dorsal and ventral cells secrete important morphogens such as BMP6 and SHH acting to regulate the growth and fate of developing spinal cord stem cells. The role and contribution of these dorsal and ventral cells to the maintenance of stem cells during aging of the human adult spinal cord niche remains to be elucidated.

While this work was in progress, a single cell RNA seq analysis was performed on seven lumbar spinal cords with some of the cases included in the present article and aged 50-80 years old. Results are released as a preprint manuscript and an associated browsable interface^22^. Using this gene expression database, we found that 6/7 of the genes we explored in our study by IHC, namely Arx, FoxJ1, Msx1, Pax6, Rfx2 and Sox2 were well expressed in ependymal cells at the single cell RNA level (fig. S3). FoxA2 expression was not detected neither in ependymal cells, nor in the entire spinal cord, maybe as the result of its expression by only a few cells located in the ventral central canal (fig. 3). These independent data clearly support our finding of the maintenance of these ependymal cells during human spinal cord aging.

In conclusion, recent work in mice, uncovered that ependymal cells are in a permissive chromatin state that enables the unfolding of a normally latent gene expression program for oligodendrogenesis which could be used to replace large numbers of lost oligodendrocytes in the injured mouse spinal cord^23^. In that spirit the maintenance of ependymal cells expressing stem cell markers throughout life in humans suggests that, theoretically, these cells could be considered as a potential source to be used for the regeneration of spinal cord lesions such as trauma or degenerative diseases. Using endogenous stem cells is however not an any task and we are far from clinical applications. Further work is needed to establish whether aged ependymal cells maintained the proliferation and differentiation capacities observed in young donors. The present work constitutes a new step toward this goal.

## Acknowledgements

We thank the French Agency of biomedicine and all Montpellier Biocampus facilities for help (RHEM) and excellent technical work. We also warmly thank all organ donors and their family as well as nurses and all medical staff from the Unit of Donation and Transplantation (Montpellier hospital). We thank Dr Ariel Levine (NIH, USA) for permission to use screen capture from the Human Spinal Cord snRNAseq Viewer (https://vmenon.shinyapps.io/humanspinalcord/)

## Supplementary material

Figure S1-3

Table S1-4

## Figures legends

**Figure S1:**
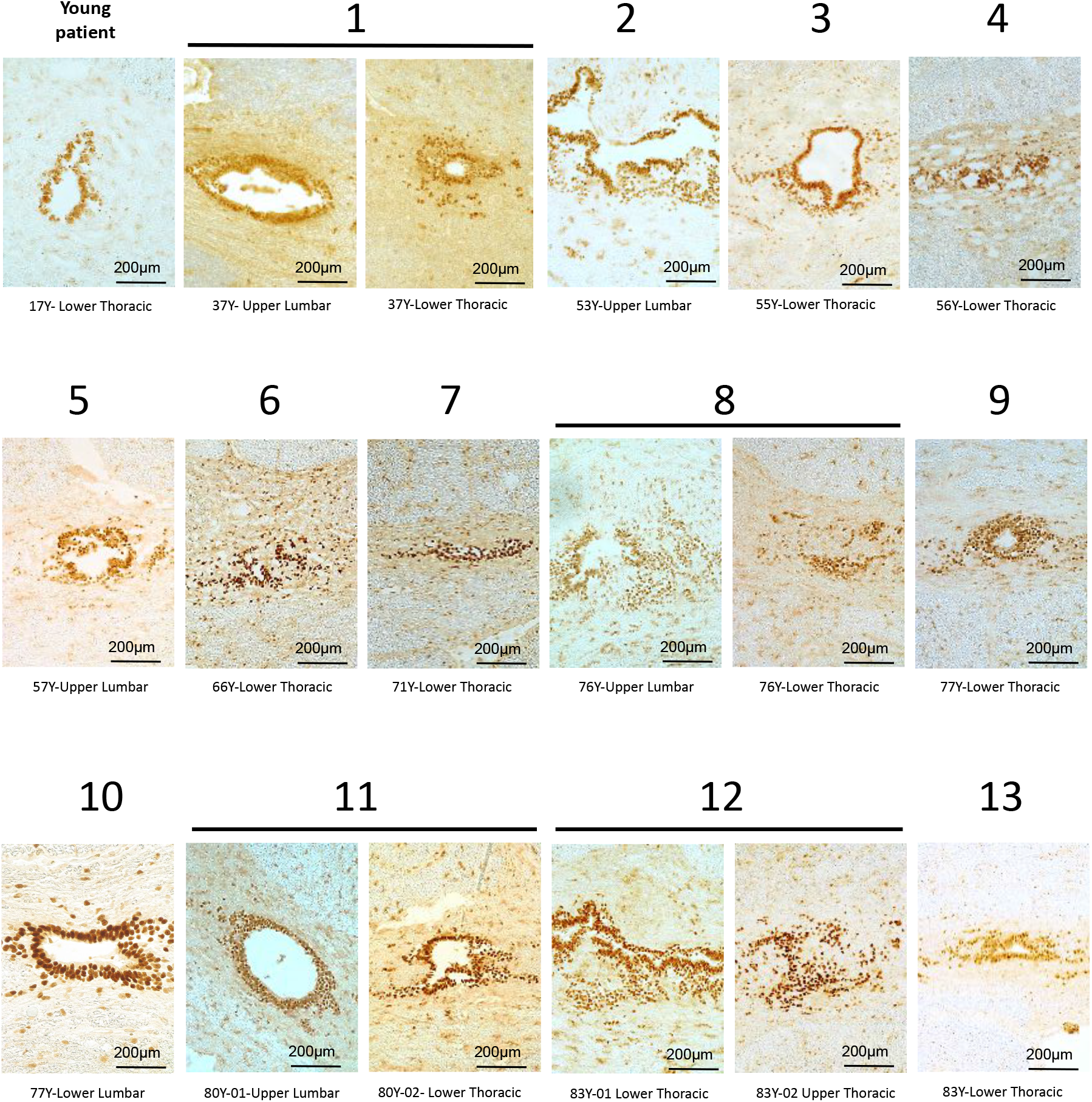
Maintenance of FOXJ1^+^ cells with aging. The series of images shows FOXJ1-stained sections in donors aged 37 to 83 years. Staining for FOXJ1 in a young donor (17 y. old), already studied in ^7^ is shown for comparison.

**Figure S2:**
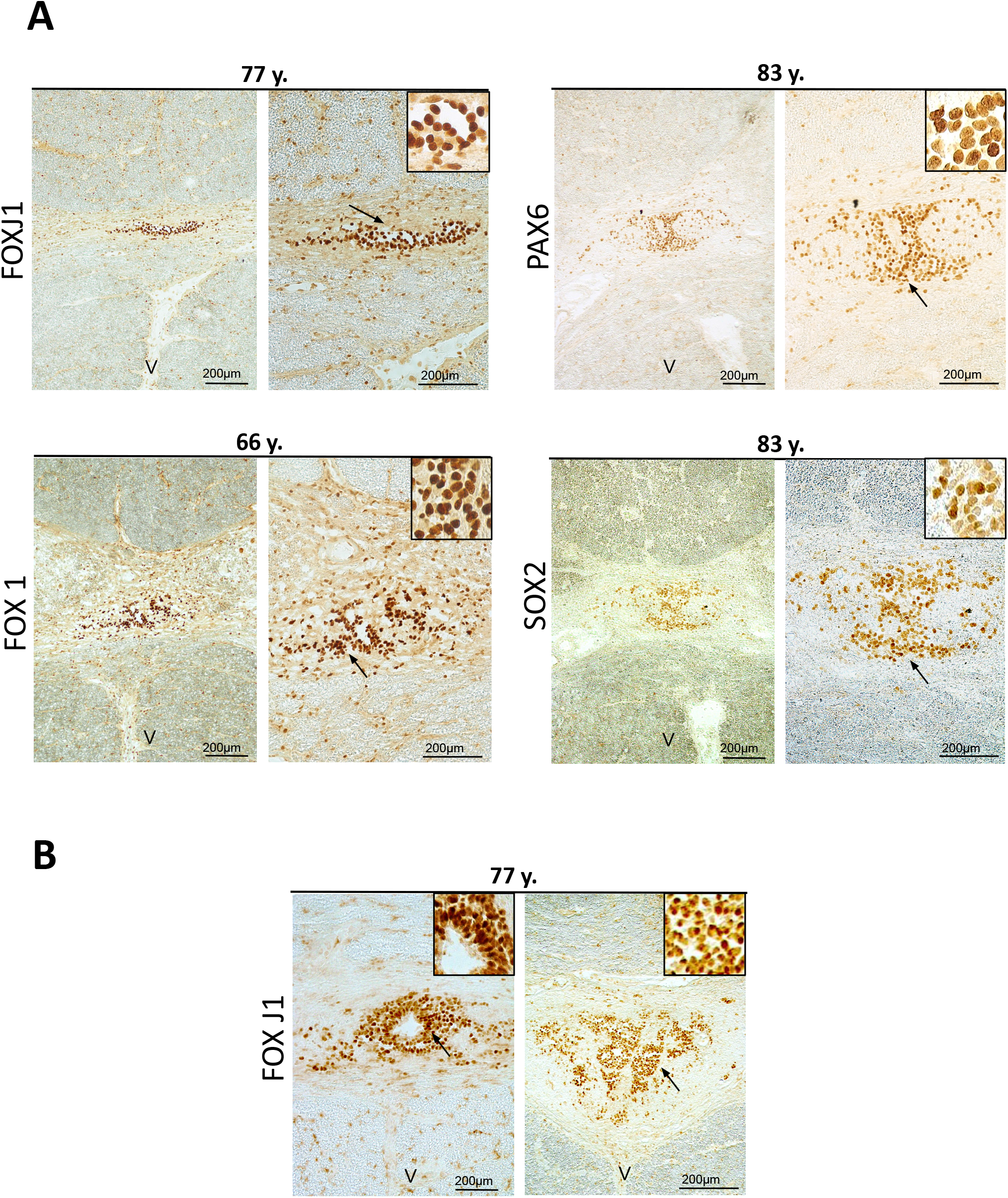
Maintenance of FOXJ1^+^, PAX6^+^, SOX2^+^ cells in stenosis. ***A***. Immunohistochemistry for indicated proteins. Black arrows indicate the magnified zone presented in the inset. ***B:*** Immunohistochemistry for FOXJ1 in one donor showing stained cells in sections with an open (left) or a closed (right) central canal.

**Figure S3:**
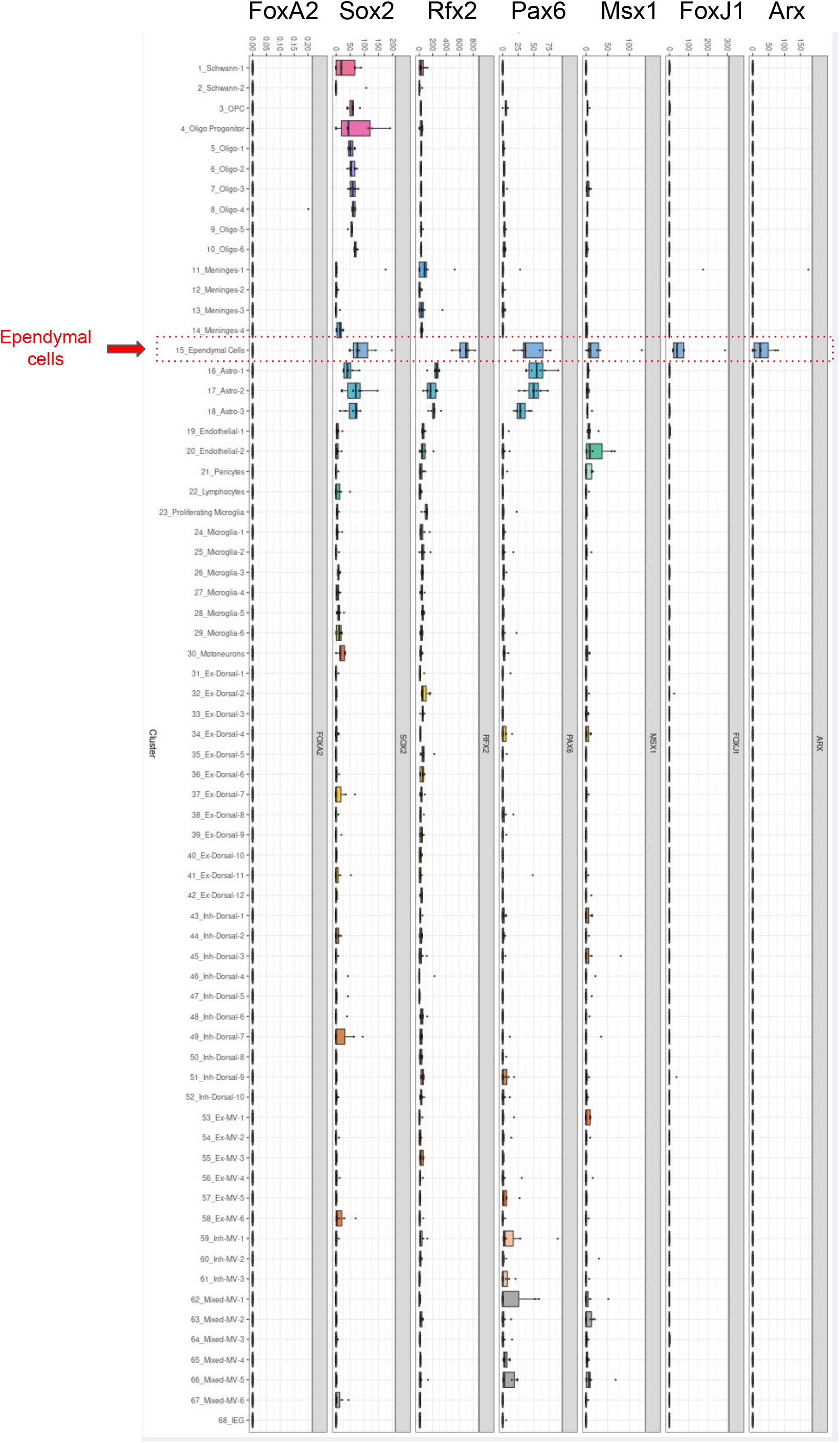
single cells RNA seq of aged spinal cords. Image downloaded from the Human Spinal Cord snRNAseq Viewer (https://vmenon.shinyapps.io/humanspinalcord/) associated with this pre-print article (Yadav, Matson, et al. 2022). Histograms show the RNA expression level in seven lumbar spinal cords of genes studied by IHC in the present article. Sixty eight cell clusters were identified by Yadav, Matson, et al. 2022. The ependymal cell cluster is indicated by the red dotted box.

