## Supplemental Table S1-S4 for "Persistence of FoxJ1^+^ Pax6^+^ Sox2^+^ ependymal cells throughout life in the human spinal cord"

Table S1 : Patient description

| Donors number | Age of donor | sex | Death cause | Elapsed time between brain death and vascular clamping | Elapsed time between vascular clamping and extraction | Elapsed time between brain death and extraction |
|---|---|---|---|---|---|---|
| 1 | 62 | M | Head trauma | 9h50 | 3h45 | 13h35 |
| 2 | 62 | M | Suicide by gunshot | 14h50 | 4h45 | 19h35 |
| 3 | 83 | M | Stroke | 7h55 | 4h45 | 12h40 |
| 4 | 77 | F | Stroke | 21h30 | 2h40 | 24h10 |
| 5 | 83 | F | posterior fossa hematoma | 22h35 | 4h15 | 26h50 |
| 6 | 53 | M | Stroke | 21h35 | 3h20 | 24h50 |
| 7 | 76 | F | Stroke | 19h20 | 3h40 | 23h00 |
| 8 | 76 | M | Stroke | 4h10 | 3h40 | 7h50 |
| 9 | 77 | M | Stroke | 2h50 | 03h25 | 06h10 |
| 10 | 80 | F | Stroke | 14h50 | 2h15 | 17h40 |
| 11 | 57 | M | Stroke | 18h00 | 3h00 | 21h05 |
| 12 | 68 | M | Aneurysm rupture | 9h26 | 3h05 | 12h30 |
| 13 | 56 | F | Aneurysm rupture | 8h25 | 2h30 | 10h55 |
| 14 | 71 | M | Stroke | 8h15 | 2h40 | 10h50 |
| 15 | 55 | F | tetraventricular hemorrhage | 15h40 | 03h50 | 19h30 |
| 16 | 66 | M | Stroke | 17h50 | 3h15 | 21h50 |
| 17 | 37 | M | Aneurysm rupture | 22h45 | 4h05 | 26h45 |

### Table S2: Antibodies

| Antibody | Species | references | Suppliers | Dilution |
|---|---|---|---|---|
| **ARX** | Polyclonal Sheep IgG | AF7068 | Biotechne R&D systems | 1:500 |
| **FOX A2** | Goat Polyclonal IgG | AF2400 | Bio-techne R&D systems | 1:500 |
| **FOX J1** | Mouse IgG1- Monoclonal Antibody (2A5) | 14-9965-82 | Bioscience™ Invitrogen | 1 :300 |
| **MSX 1** | Polyclonal Goat IgG | AF5045 | Biotechne R&D systems | 1:500 |
| **PAX 6** | Rabbit Polyclonal | 901301 | BioLegend | 1:600 |
| **RFX 2** | Rabbit Polyclonal | HPA048969 | SIGMA-MERCK | 1:500 |
| **SOX 2** | Rabbit Polyclonal | 14962 | Cell Signaling | 1:500 |

### Table S3: State of central canal

| Donor #<br>Spinal Cord Region | Examined lenght µm (section #) | Age | Canal<br>Open/Closed |
|---|---|---|---|
| 1<br>Upper Lumbar | 140 (10 sections) | 62 | Closed |
| 2<br>Upper Lumbar | 224 (16 sections) | 62 | Closed |
| 3<br>Lower Thoracic | 588 (82 sections) | 83 | Open |
| 4<br>Lower Thoracic | 336 (24 sections) | 77 | Open/Closed |
| 5-01<br>Lower Thoracic | 6440 (460 sections) | 83 | Open |
| 5-02<br>Lower Thoracic | 336 (24 sections) | 83 | Closed |
| 6<br>Upper Lumbar | 350 (25 sections) | 53 | Open/Closed |
| 7<br>Upper Lumbar | 378 (27 sections) | 76 | Closed |
| 8<br>Lower Thoracic | 322 (23 sections) | 76 | Closed |
| 9<br>Upper Lumbar | 532 (38 sections) | 77 | Open/Closed |
| 10-01<br>Upper Lumbar | 1120 (80 sections) | 80 | Open |
| 10-02<br>Lower Thoracic | 42 (3 sections) | 80 | Open |
| 11<br>Lower Thoracic | 742 (53 sections) | 57 | Open |
| 12<br>Upper Lumbar | 378 (27 sections) | 68 | Open |
| 13<br>Lower Thoracic | 602 (43 sections) | 56 | Closed |
| 14<br>Lower Thoracic | 532 (38 sections | 71 | Closed |
| 15<br>Lower Thoracic | 42 (3 sections) | 55 | Open |
| 16<br>Lower Thoracic | 29 (2 sections) | 66 | Closed |
| 17-01<br>Upper Lumbar | 336 (24 sections) | 37 | Open |
| 17-02<br>Lower Thoracic | 84 (6 sections) | 37 | Open |

Table S4: Central Canal Stenosis

| Spinal cord | Open Canal | Closed Canal | Open/Closed Canal |
|---|---|---|---|
| Upper Lumbar 8 Samples | 3 | 3 | 2 |
| Lower Thoracic 12 Samples | 6 | 5 | 1 |
| Total | 9 | 8 | 3 |
| Pourcentage | 45% | 40% | 15% |
